# CCT3 drives Sorafenib resistance by inhibiting TFRC-mediated iron uptake in HCC

**DOI:** 10.1101/2023.12.14.571783

**Authors:** Huihui Zhu, Qiuhong Liu, Qinna Meng, Linjian Zhang, Jiaheng Lan, Danhua Zhu, Yonxia Chen, Nadire Aishan, Xiaoxi Ouyang, Sainan Zhang, Lidan Jin, Lanlan Xiao, Linbo Wang, Lanjuan Li, Feiyang Ji

## Abstract

Sorafenib is commonly utilized in the management of advanced hepatocellular carcinoma (HCC). However, its efficacy in extending patients’ survival is hindered by the development of drug resistance. By employing protein posttranslational modification (PTM) omics, including acetylome, phosphoproteome, and ubiquitinome, in conjunction with genome-wide CRISPR/Cas9 knockout library screening, we have successfully identified chaperonin containing TCP1 subunit 3 (CCT3) as a key factor contributing to Sorafenib resistance. Furthermore, we observed a reduction in the ubiquitination of CCT3 at lysine 21 (K21) subsequent to Sorafenib treatment. This study provides evidence that CCT3 hinders the recycling of transferrin receptor protein 1 (TFRC) by interacting with alpha-actinin-4 (ACTN4), which is influenced by K6-linked ubiquitination on K21. Depleting CCT3 increased the susceptibility of cells to Sorafenib-induced ferroptosis, while reintroducing CCT3 through transfection restored resistance to ferroptosis. Additionally, impairing ACTN4 or TFRC depletion compromised CCT3’s ability to inhibit Sorafenib-induced ferroptosis. In summary, targeting CCT3 presents a potential strategy for overcoming Sorafenib resistance in HCC.

## Introduction

Liver cancer holds the seventh position globally among all cancer types, and it stands as the second leading cause of cancer-related deaths(Sung et al., 2021). Among primary liver cancers, hepatocellular carcinoma (HCC) is the most prevalent. In the initial stages of HCC, surgical intervention remains the prevailing treatment modality. However, approximately 50% of advanced HCC cases necessitate Sorafenib and local ablation as the primary therapeutic approach(Forner et al., 2018; Llovet et al., 2016). Sorafenib, an orally administered multi-target tyrosine kinase inhibitor, effectively impedes the proliferation and angiogenesis of tumor cells by targeting vascular endothelial growth factor receptor (VEGFR), platelet-derived growth factor receptor (PDGFR), Raf family kinases, among others. Consequently, the survival rates of HCC patients experience significant improvement(Liu et al., 2006; Llovet et al., 2008). Despite its effectiveness in treating HCC, Sorafenib’s impact on patients’ median overall survival is limited, extending it by merely three months(Llovet et al., 2008). The emergence of drug resistance poses a significant challenge to the efficacy of Sorafenib treatment in HCC patients. Enhancing our comprehension of the molecular mechanisms underlying Sorafenib resistance in HCC holds the potential to enhance disease prognosis and facilitate the identification of therapeutic targets capable of circumventing chemoresistance.

Epigenetics, transport processes, tumor microenvironment, and regulated cell death have been identified as potential triggers and contributors to the development of Sorafenib resistance(Tang et al., 2020). In the realm of chemotherapy resistance mechanisms, there has been a growing focus on ferroptosis, which exhibits unique characteristics in terms of morphology, biochemistry, and genetics compared to apoptosis, autophagy, necrosis, and other forms of cell death(Dixon et al., 2012). In the process of ferroptosis, there is a significant accumulation of iron and lipid peroxidation. The regulatory mechanisms primarily involve Xc-system, glutathione peroxidase 4(GPX4) activity, and reactive oxygen species (ROS) production(Li et al., 2020). Sorafenib, unlike other tyrosine kinase inhibitors, has the ability to induce ferroptosis in various cancer cell lines(Lachaier et al., 2014). Research has indicated that ferroptosis is linked to Sorafenib resistance in HCC. For instance, the depletion of intracellular iron stores using deferoxamine (DFO) has been shown to protect HCC cells from the cytotoxic effects of Sorafenib(Louandre et al., 2013). However, further investigation is required to elucidate the molecular mechanism of ferroptosis in Sorafenib resistance.

CCT3, a subunit of the chaperonin containing TCP1 (CCT) complex, is a constituent of the TCP1 ring complex (TRiC) family, comprising eight subunits (CCT1-CCT8). The CCT complex serves as a molecular chaperone essential for the proper folding of actin and tubulin, key components of cytoskeletal microfilaments and microtubules. Additionally, the individual subunits of the CCT complex have been observed to possess independent functionalities(Grantham, 2020). Increased expression of CCT3 has been associated with unfavorable prognoses in patients with HCC, while its depletion has been shown to induce apoptosis and impede the proliferation of liver cancer cells(Cui et al., 2015; Hou et al., 2019). Furthermore, existing research has demonstrated that CCT3 exerts inhibitory effects on apoptosis, while simultaneously promoting cell proliferation and metastasis in various malignant tumors, including breast cancer, melanoma, and lung cancer(Chen et al., 2022a; Liu et al., 2022; Temiz et al., 2021). However, the potential contribution of CCT3 to Sorafenib resistance in HCC remains uncertain, although it has been observed to induce cisplatin resistance in lung adenocarcinomas through its targeting of JAK2/STAT3(Danni et al., 2021). Moreover, the inhibition of ferroptosis by CCT3 has been implicated in the development of lung cancer, while its presence can lead to the inhibition of lipid metabolism and the promotion of lipid accumulation in liver cancer(Sondergaard et al., 2022; Wang et al., 2022).

Sorafenib has the potential to induce signal transduction and posttranslational modification (PTM) alterations in hepatoma cells. Consequently, the identification of proteins exhibiting significant PTM changes can serve as a means to screen for molecules associated with Sorafenib resistance. Mass spectrometry-based protein PTM omics enables the simultaneous identification and quantification of numerous protein PTMs, offering advantages such as high throughput, precision, and the ability to directly localize modifications to specific amino acid residues. As a result, this methodology has gained increasing prominence in cancer research in recent years. Wang et al. have identified phosphorylated PTPN11 and PLCG1 as potential mediators of oncogenic pathway activation in human glioblastoma (Wang et al., 2021b). Similarly, another study utilizing this approach has demonstrated the critical role of SYVN1 in the progression of HCC metastasis(Ji et al., 2021). Genome-wide CRISPR/Cas9 screening involves the utilization of CRISPR/Cas9 technology to construct a comprehensive gene mutation library of a particular species. This library is then subjected to functional screening, PCR amplification, and deep sequencing to identify genes associated with a specific phenotype. Lai Wei and colleagues employed this approach to conduct a screening of 986 genes implicated in Sorafenib resistance in 97L cells(Wei et al., 2019). Huang and colleagues discovered that the absence of DUSP4 leads to the development of lenvatinib resistance in HCC(Huang et al., 2022). The integration of PTM omics and genome-wide Cas9 screening enables a comprehensive identification of proteins associated with specific biological processes, encompassing signaling pathways and phenotypes. This methodology significantly enhances the precision of screening and provides valuable insights for subsequent mechanistic investigations.

## Results

### Integrating PTM omics and CRISPR/Cas9 knockout screening identified CCT3 as a resistance gene to Sorafenib

In this study, we conducted PTM omics analysis to quantify alterations in protein phosphorylation, acetylation, and ubiquitination following Sorafenib treatment in HCC cells (Fig 1A, Supplementary Table 1). The genome-wide CRISPR/Cas9 knockout library screening was previously carried out by Lai et al(Wei et al., 2019). The experimental parameters for the selection of 97L cells and the administration of 7uM Sorafenib in PTM omics were adopted from the CRISPR/Cas9 knockout library screening to ensure the congruity of the findings between the two approaches. Using PTM omics, we successfully detected a total of 224 acetylated modified proteins, 168 phosphorylated modified proteins, and 442 ubiquitinated modified proteins in 97L cells. Interestingly, 11 proteins exhibited all three types of PTMs (Fig 1B). Upon treatment with Sorafenib, we observed alterations in the PTMs of certain proteins, primarily an increase in their abundance (Fig S1A). Furthermore, our analysis of gene ontology (GO) terms enrichment indicated that these proteins were associated with membrane raft, chaperonin-containing T-complex, and receptor tyrosine kinase binding (Fig S1B-C). Among the list of 14 proteins whose PTM was significantly altered following treatment with Sorafenib, it was found that only CCT3 had an impact on Sorafenib resistance as determined through screening with a genome-wide CRISPR/Cas9 knockout library (Fig 1C). Specifically, the ubiquitination of lysine residue 21 (K21) in CCT3 was notably reduced after Sorafenib treatment, particularly at the 30- minute time point, indicating the crucial involvement of CCT3 in the signaling pathway of 97L cells in response to Sorafenib (Fig 1D-E, S1D). Furthermore, the presence of CCT3 targeting single guide RNA (sgRNA) was noticeably diminished in Sorafenib-treated 97L cells, suggesting that the depletion of CCT3 may enhance sensitivity to Sorafenib (Fig 1F). The findings from the aforementioned methodologies indicate that CCT3 exhibits responsiveness to Sorafenib stimuli and can effectively inhibit its activity. Additionally, Sorafenib disrupts global phosphorylation, acetylation, and ubiquitination, suggesting their involvement in cellular responses (Fig S1E). The iceLogo analysis further revealed conserved patterns of peptides with notable PTM alterations (Fig S1F).

**Fig. 1.**
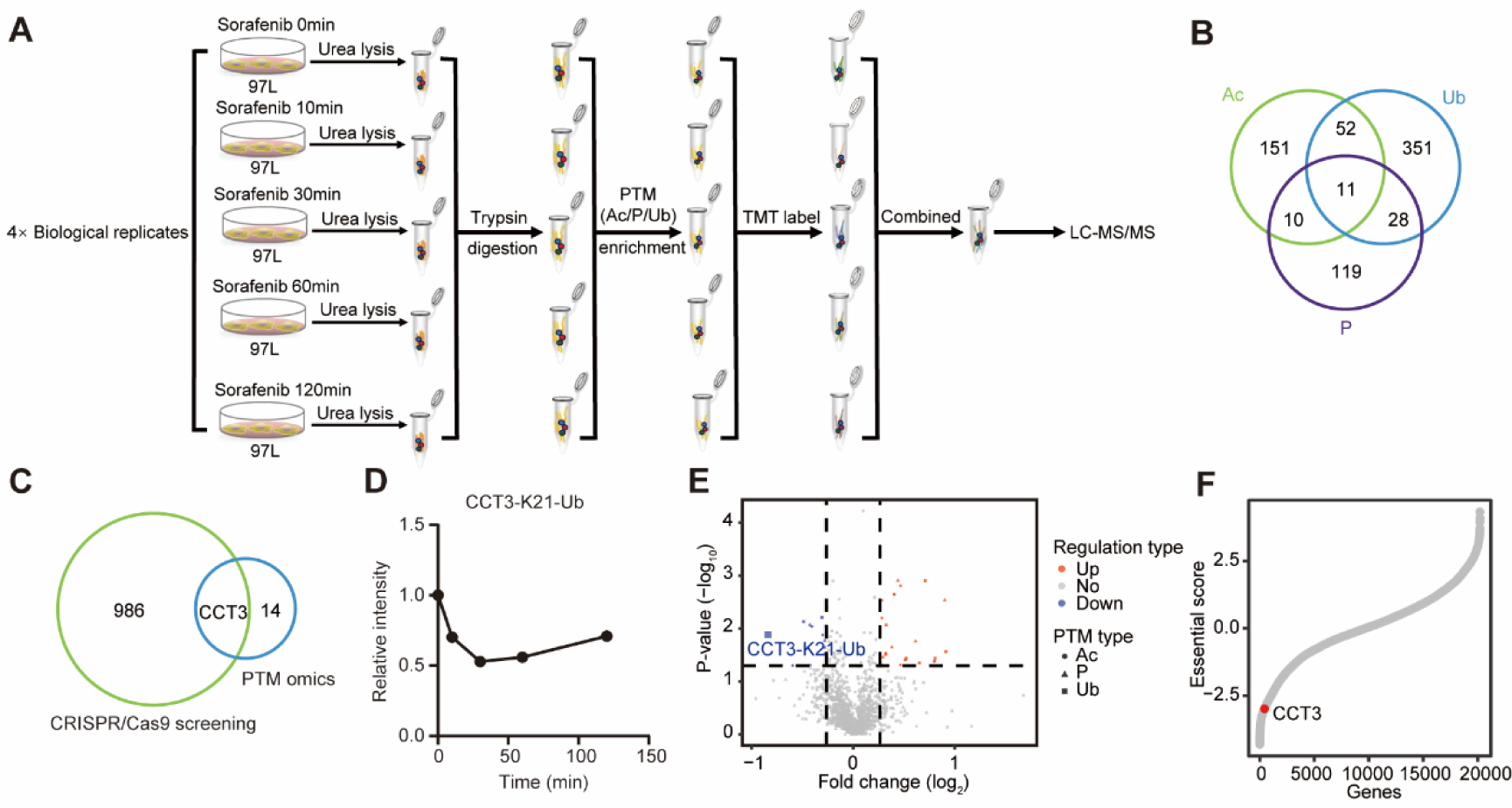
Combined PTM omics and genome-wide CRISPR/Cas9 knockout identified CCT3 as an important gene for Sorafenib sensitivity. A. Flow diagram depicting the posttranslational modification (PTM) omics of acetylation (Ac), phosphorylation (P), and ubiquitination (Ub) in 97L cells treated with Sorafenib was presented. **B** Venn diagram showing the overlap of identified proteins between three posttranslational modifications. **C** Venn diagram displaying the overlap of proteins across PTM omics screening and genome-wide CRISPR/Cas9 knockout screening. **D** The line chart shows the change of the ubiquitination of CCT3 at lysine 21 (CCT3-K21-Ub) in PTM omics after Sorafenib treatment. **E** Volcano plot showing the regulated PTM sites in 97L cells treated with Sorafenib for 1 h. **F** The essential score of CCT3 targeting sgRNAs (red dot) in genome-wide CRISPR/Cas9 knockout screening. Negative essential scores represent negative selection, and positive essential scores represent positive selection.

### CCT3 promotes Sorafenib resistance in HCC cells

In order to validate the findings obtained through the combination of two screening methods, the transcriptomic dataset GSE109211 of HCC patients treated with Sorafenib was obtained from the Gene Expression Omnibus database. The analysis revealed a negative correlation between the expression level of CCT3 in HCC and the efficacy of Sorafenib (Fig 2A). Additionally, to further investigate the impact of CCT3 on the susceptibility of HCC to Sorafenib, two small interfering RNA (siRNA) sequences targeting CCT3 were designed and utilized to suppress CCT3 expression in two HCC cell lines, namely 97L and LM3. The knockdown efficiency of two CCT3 siRNA molecules was further confirmed through western blotting analysis in both 97L and LM3 cells (Fig 2B, S2A). The knockdown of CCT3 resulted in increased sensitivity of 97L and LM3 cells to Sorafenib-induced growth inhibition (Fig 2C). Notably, the rescue experiment demonstrated that the overexpression of CCT3 mRNA reversed the effect of CCT3 siRNA on Sorafenib sensitivity (Fig 2D). Additionally, the knockdown of CCT3 significantly impaired the ability of 97L and LM3 cells to form clones (Fig 2E-F, S2B-C). Using flow cytometry, we observed a statistically significant increase in cell death following CCT3 knockdown in Sorafenib-treated cells (Fig 2G-H, S2D-E). These results suggest that inhibiting CCT3 enhances the susceptibility of HCC cells to Sorafenib therapy.

**Fig. 2.**
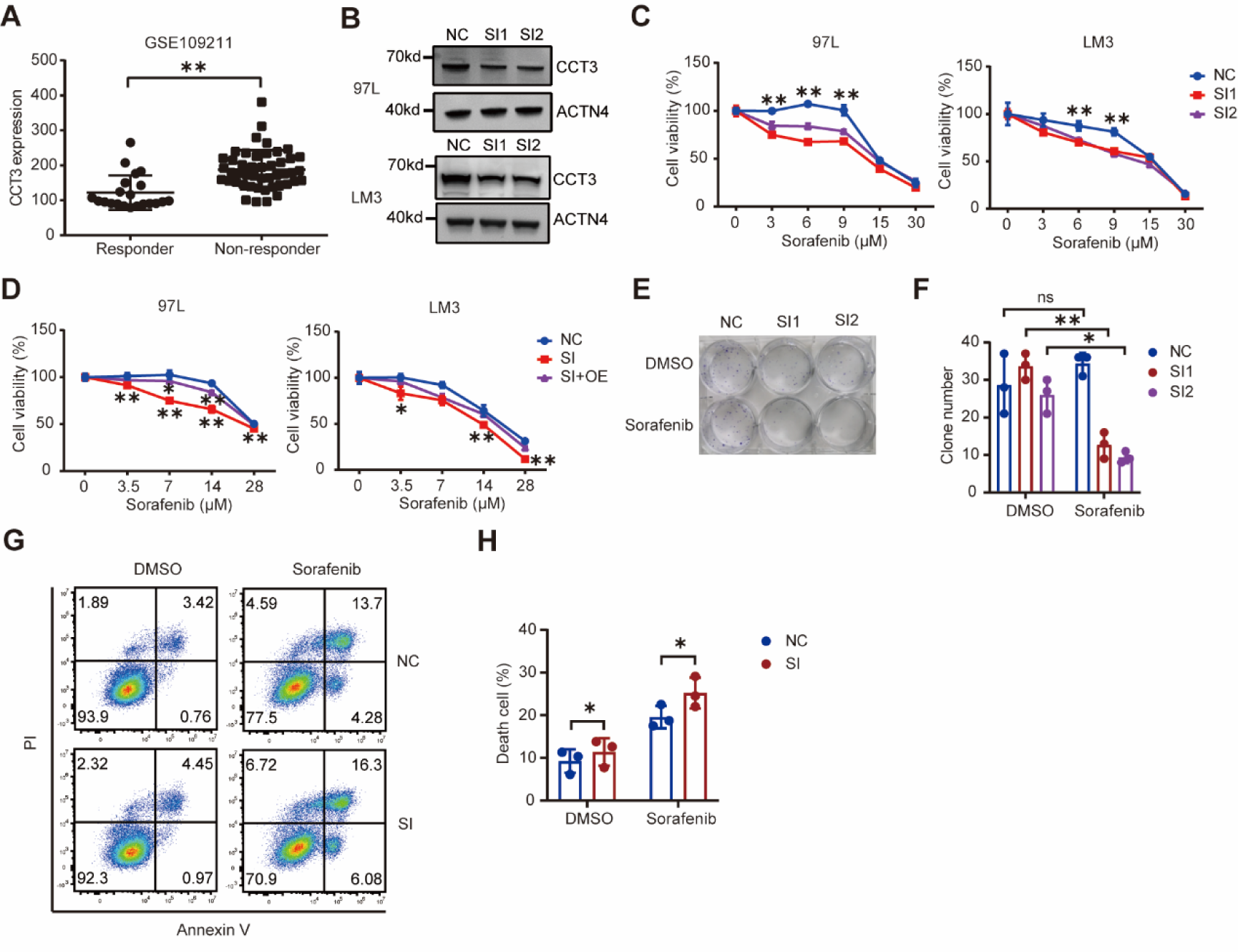
Inhibiting CCT3 sensitized HCC cells to Sorafenib. A. The expression level of CCT3 in Sorafenib response or non-response hepatocellular carcinoma from GEO database. **B** Western blot analysis of the indicated proteins in cells treated with negative control siRNA (NC) and two CCT3 siRNA (SI1, SI2). **C** Analysis of cell viability in control and CCT3-knockdown 97L and LM3 cells following treatment with Sorafenib for 48 h. **D** Cell viability of control, CCT3- knockdown (SI), and CCT3-overexpression (OE) after CCT3-knockdown cells following treatment with Sorafenib for 24 h. **E** Clone formation capacity of control and CCT3-knockdown 97L cells following treatment with DMSO or 2 µM Sorafenib. **F** Statistics histogram for clone assays from three biological replicates. **G** Apoptosis of indicted 97L cells treated with DMSO or 7 µM Sorafenib for 24 h were performed using Annexin V/PI staining. **H** Histogram showing the statistical results of apoptosis from three biological replicates. Each cell viability data is representative of three biological replicates.

### CCT3 promotes Sorafenib resistance by inhibiting ferroptosis

To investigate the underlying mechanism through which CCT3 facilitates resistance to Sorafenib in HCC, a comprehensive analysis of cellular proteomics was conducted to identify alterations in protein expression in 97L and LM3 cells following treatment with Sorafenib and inhibition of CCT3 expression (Fig 3A, Supplementary Table 2). The results of our study demonstrated significant changes in protein expression in 97L and LM3 cells upon Sorafenib treatment and CCT3 inhibition (Fig S3A). The Kyoto Encyclopedia of Genes and Genomes (KEGG) pathway analysis indicated that the proteins that exhibited differential expression after inhibiting CCT3 were implicated in various cellular processes, including glutathione metabolism, mitophagy, and endocytosis, which are associated with intracellular iron transport and ferroptosis (Fig 3B). Additionally, gene set enrichment analysis (GSEA) confirmed the involvement of these proteins in ferroptosis (Fig 3C). The effectiveness of CCT3 knockdown was also confirmed through GO analysis, which revealed enrichment of the chaperonin-containing T-complex pathway (Fig S3B). The analysis conducted using KEGG and GSEA also revealed the involvement of ferroptosis, reactive oxygen species, and oxidative phosphorylation in the response to Sorafenib (Fig S3C-D). The collective findings from the proteomics study suggest that the inhibition of CCT3 enhances Sorafenib sensitivity, potentially through the induction of ferroptosis. To confirm this hypothesis, we employed 3-Methyladenine (3-MA), Necrostatin-1 (Nec-1), Z-VAD-FMK, Deferoxamine (DFO), and Ferrostatin-1 (Fer-1) to inhibit autophagy, necrosis, apoptosis, and ferroptosis, respectively. Our results demonstrated that only the ferroptosis inhibitors DFO and Fer-1 weakened the impact of CCT3 on Sorafenib sensitivity, while the other inhibitors of cell death pathways did not (Fig 3D). In order to investigate the central events of ferroptosis, namely iron accumulation and lipid peroxidation, we conducted further analysis. Utilizing FerroOrange, a probe for detecting ferro-ions, we observed that the downregulation of CCT3 resulted in enhanced ferrous ion accumulation in Sorafenib-treated 97L and LM3 cells (Fig 3E-F, S3E-F). Additionally, the knockdown of CCT3 promoted lipid peroxidation in these cells (Fig 3G-H, S3G-H). Consistent with expectations, our findings demonstrated that the reduction of CCT3 levels led to a decrease in intracellular glutathione levels (Fig 3I, S3I). The findings from transmission electron microscopy demonstrated that the reduction of CCT3 expression resulted in a decrease in mitochondrial volume and an increase in membrane density in 97L and LM3 cells (Fig 3J, S3J). Notably, the rescue experiment revealed that the overexpression of CCT3 mitigated the impact of CCT3 expression interference on the accumulation of ferrous ions in 97L and LM3 cells treated with Sorafenib (Fig 3K-L, S3K-L). In conclusion, CCT3 functions as an inhibitor of Sorafenib-induced ferroptosis in HCC cells.

**Fig. 3.**
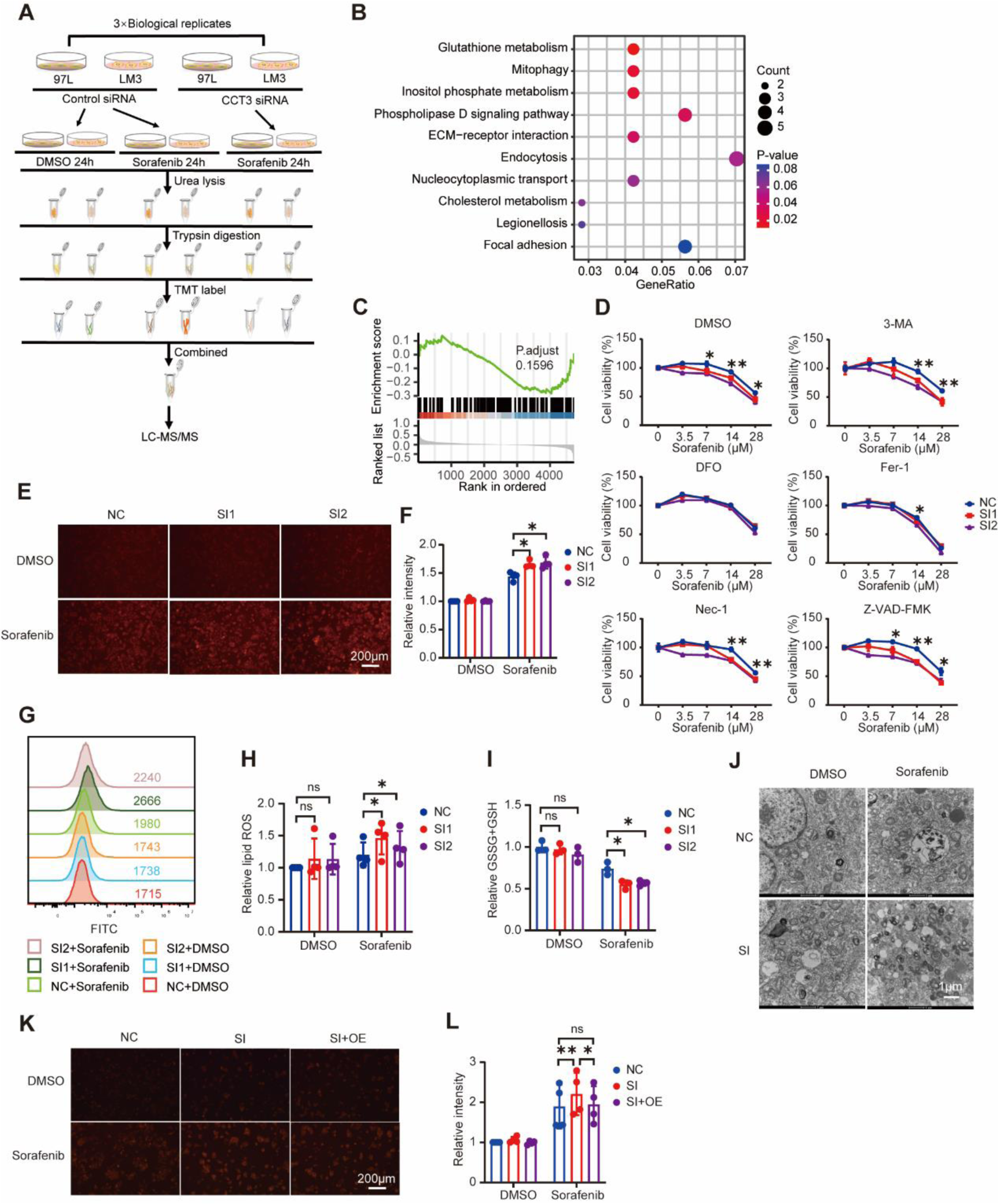
Inhibition of CCT3 promotes ferroptosis in Sorafenib-treated HCC cells. A. Flow chart of the whole cell proteomic for 97L and LM3 cells treated with Sorafenib and CCT3 siRNA. **B** KEGG pathway enrichment analysis of differentially expressed protein between control and CCT3-knockdown 97L cells. **C** GSEA ferroptosis pathway enrichment analysis of differentially expressed protein between control and CCT3-knockdown 97L cells. **D** Cell viability of control and CCT3-knockdown 97L cells treated with Sorafenib and indicated cell death inhibitors for 24 h. **E-F** The accumulation of iron in control and CCT3-knockdown 97L cells were detected using FerroOrange after treated with DMSO or Sorafenib (14 µM) for 12 h. Histogram showing the statistical results of iron accumulation from four biological replicates. **G-H** 97L cells were treated with DMSO or Sorafenib (14 µM) for 24 h, and then lipid hydroperoxides were measured. Statistics for the median Fluorescence intensity of oxidized BODIPY dyes were carried out on four biological replicates. **I** 97L cells were treated with Sorafenib at 14 µM for 12 h, and the intracellular glutathione (GSH) level were assayed. **J** Transmission electron microscopy was used to examine the morphology of indicated 97L cells after 24 hours of treatment with Sorafenib (14 µM). **K-L** Indicated 97L cells were treated with Sorafenib (14 µM) for 12 h, and then iron accumulation were measured and statistical analysis.

### CCT3 interacts with ACTN4

To investigate the inhibitory mechanism of ferroptosis by CCT3, we employed co-immunoprecipitation in conjunction with mass spectrometry to identify potential protein interactions with CCT3 (Figure 4A). The proteins that exhibited a greater pull-down by CCT3 antibodies compared to the IgG isotype control were compiled (Figure 4B). Among these proteins, ACTN4 displayed the highest protein abundance. Furthermore, the presence of CCT3 and other subunits of the TRiC, namely CCT4, CCT5, and CCT8, were also detected, indicating the successful implementation of our methodology. Notably, we observed a significant enhancement in the binding between CCT3 and ACTN4 upon Sorafenib treatment (Figure S4A). Using Pymol, the identification and classification of all functional residues were conducted based on their interactions with other proteins. Specifically, the residues LYS78 of CCT3 and ASN73 of ACTN4 were found to form hydrogen bonds, indicating a strong interaction. The CCT3-ACTN4 interaction score was determined to be -557, which is considered favorable (Fig 4C). To further validate the interaction between CCT3 and ACTN4, endogenous co-immunoprecipitation experiments were performed in 97L and LM3 cell lines. The results demonstrated that both anti-CCT3 antibodies and anti-ACTN4 antibodies were able to simultaneously pull down CCT3 and ACTN4 (Fig 4D-E, S4B-C). In addition, exogenous co-immunoprecipitation was performed in 293T cells, revealing that the Flag-CCT3 and Myc- ACTN4 proteins were simultaneously pulled down by the anti-Flag antibody and anti-Myc antibody (Fig 4F, S4D). Colocalization of CCT3 and ACTN4 was also observed through confocal laser scanning microscopy (Fig 4G). Consequently, it can be concluded that CCT3 interacts with ACTN4, and the administration of Sorafenib may enhance this interaction.

**Fig. 4.**
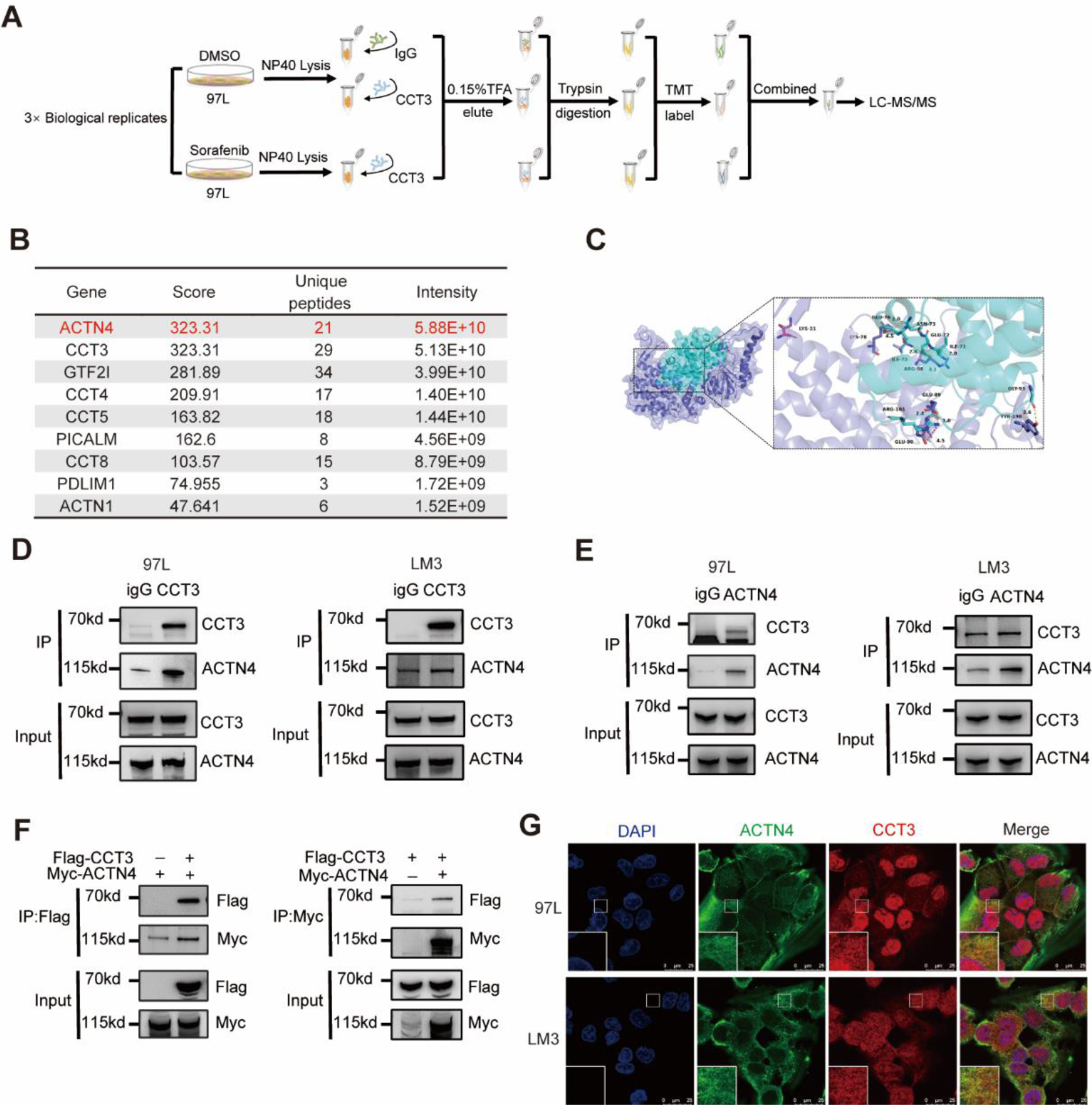
ACTN4 is a interacting protein of CCT3. A. Flow chart showing the combined use of immunoprecipitation and proteomics to identify the partners of CCT3. **B** List of proteins that may interact with CCT3, sorted by intensity. **C** Molecular docking of the CCT3 (slate cartoon) and ACTN4 (cyan cartoon) interaction complex, with corresponding- colored stick structures representing the binding sites. **D** Analysis of CCT3-binding protein in 97L and LM3 cells by co- immunoprecipitation. **E** Analysis of ACTN4-binding protein in 97L and LM3 cells by co-immunoprecipitation. **F** Analysis of CCT3 and ACTN4 binding protein in 293T cells by co-immunoprecipitation. **G** Immunofluorescence staining showing the distribution of CCT3 (red), ACTN4 (green) and DAPI (blue) in indicated cells.

### The CCT3/ACTN4 axis inhibits ferroptosis by impeding TFRC recycling

ACTN4 has been previously documented as essential for the efficient recycling of TFRC, which is responsible for maintaining sufficient TFRC levels on the cell membrane to support iron endocytosis (Yan et al., 2005). Consequently, we postulated that CCT3 may impede Sorafenib-induced ferroptosis by obstructing iron uptake mediated by ACTN4. To substantiate this conjecture, we initially assessed the impact of Sorafenib and CCT3 knockdown on the expression of proteins associated with ferroptosis. Western blot analysis revealed a reduction in GPX4 levels in 97L and LM3 cells following Sorafenib treatment, indicating the induction of ferroptosis by Sorafenib. Furthermore, the downregulation of CCT3 did not exhibit any impact on the levels of ACTN4, Ferritin, and TFRC, indicating that CCT3 may potentially interact with ACTN4 to influence its capacity to sustain TFRC recycling, rather than influencing its expression or degradation (Fig 5A, S4E-F). To validate the influence of CCT3 on iron endocytosis, 97L and LM3 cells were cultivated in serum-free medium supplemented with FITC-transferrin for a duration of 15 minutes, followed by removal and fixation at 4% polyformaldehyde, and subsequent analysis of fluorescence intensity using flow cytometry. Unsurprisingly, the fluorescence intensity of FITC in cells with CCT3 knockdown exhibited a significant decrease compared to that in control cells (Fig 5B-C). The intracellular level of transferrin is known to be influenced by various processes such as endocytosis, recycling, and lysosomal degradation(Maxfield and McGraw, 2004; Wang et al., 2021a). Consequently, we investigated the specific impact of CCT3 on these processes by utilizing markers for early endosomes (EEA1), recycling endosomes (RAB4), and late endosomes (RAB7)(Eggers et al., 2009; Gryaznova et al., 2018; Langemeyer et al., 2018). Through immunofluorescent confocal laser microscopy, it was observed that CCT3 co-localized with RAB4, while no co-localization was observed with EEA1 or RAB7 in 97L and LM3 cells (Fig 5D). ACTN4 was found to co- locate with EEA1 and RAB4, while TFRC was found to co-locate with EEA1, RAB4, and RAB7, which is consistent with previous research (Fig S5A-B). To further substantiate the impact of CCT3 on the recycling process of TFRC, we examined the alterations in the co-locational Pearson correlation coefficients between TFRC and endosome markers following CCT3 knockdown. The findings revealed a significant decrease in the co-location between TFRC and RAB4 after CCT3 knockdown, a slight decrease in the co-location between TFRC and EEA1, and no change in the co-location between TFRC and RAB7 (Fig 5E-F, S5C-F). This finding aligns with previous result indicating that the inhibition of CCT3 leads to an increased rate of TFRC recycling, ultimately resulting in a reduction of TFRC within the recycling endosome. Finally, we conducted a series of representation experiments to demonstrate that ACTN4 and TFRC are indispensable for CCT3 to perform its anti-ferroptosis function. The expression of ACTN4 and TFRC in 97L and LM3 cells was effectively silenced using siRNA (Fig 5G, S5G-I). The CCK8 assay revealed that the impact of CCT3 knockdown on Sorafenib sensitivity in HCC cells was no longer observed upon simultaneous knockdown of ACTN4 or TFRC (Fig 5H, S5J). Furthermore, the observed phenomenon of CCT3 knockdown promoting the accumulation of ferrous ions in Sorafenib-treated cells was not evident after knockdown of ACTN4 or TFRC (Fig 5I-J, S5K-L). In conclusion, the inhibition of ferroptosis by CCT3 primarily stems from its disruption of ACTN4’s role in TFRC recycling.

**Fig. 5.**
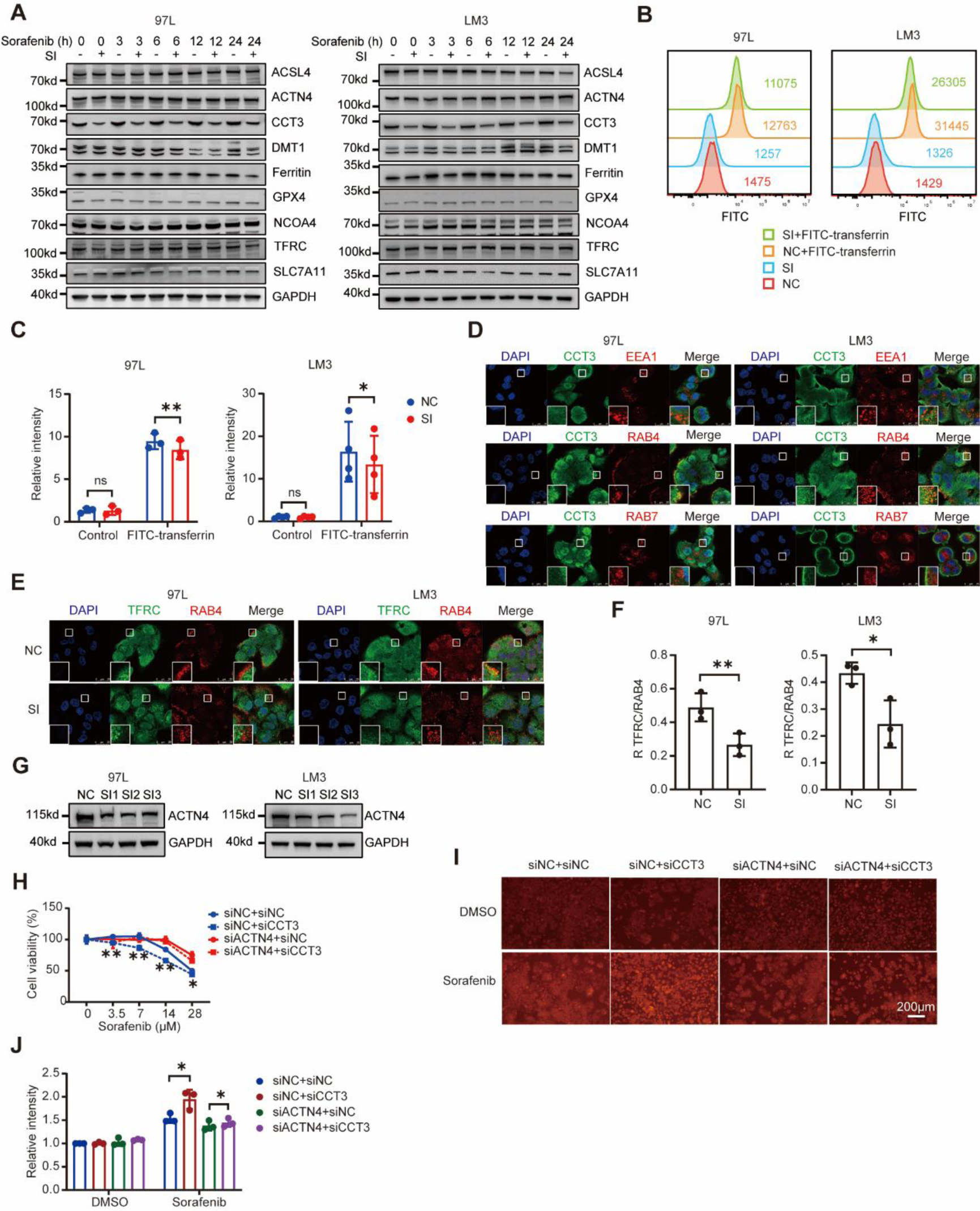
CCT3 inhibits iron endocytosis by impeding TFRC recycling through ACTN4. A. Western blot analysis of the indicated proteins in control and CCT3-knockdown 97L and LM3 cells after treated with Sorafenib (7 µM). **B** Indicated cells were incubated in fetal bovine serum free medium for 60 min and then incubated in medium contained 50 µg/ml FITC-transferrin for 15 min. Fluorescence intensity of intracellular FITC was measured by flow cytometry. **C** Histogram showing the statistical results of median FITC fluorescence intensity for indicated groups from three biological replicates. **D** Immunofluorescence staining showing the distribution of CCT3 (green), EEA1 (red), RAB4 (red), RAB7 (red) and DAPI (blue) in indicated cells. **E** Immunofluorescence staining showing the distribution of RAB4 (red), TFRC (green) and DAPI (blue) in control and CCT3-knockdown cells. **F** Pearson’s R correlation value between RAB4 and TFRC in indicated cells. **G** Western blot analysis of the knockdown efficiency of ACTN4 in 97L and LM3 cells. **H** Cell viability analysis of control (siNC), CCT3-knockdown (siCCT3), and ACTN4-knockdown (siACTN4) 97L cells following treatment with Sorafenib for 24 h. **I-J** Indicated 97L cells were treated with Sorafenib (14 µM) for 12 h, and then iron accumulation were measured. Histogram showing the statistical results from three biological replicates.

### The ferroptosis inhibition function of CCT3 requires deubiquitination at K21

In previous PTM omics analyses, it was observed that Sorafenib treatment resulted in a decrease in ubiquitination at K21 in CCT3 (Fig 1D-E). This finding suggests that ubiquitination may play a significant role in CCT3’s ability to inhibit ferroptosis. To further investigate this, a combination of immunoprecipitation and Western blotting analysis was conducted, which confirmed that Sorafenib treatment reduced the ubiquitination of CCT3 in HCC cells (Fig 6A, S6A). Additionally, it was observed that Sorafenib did not decrease the ubiquitination of CCT3 when the K21 residue was mutated to arginine (K21R), indicating that Sorafenib primarily affects the ubiquitination of CCT3 at K21 (Fig 6B, S6B). Using lysine-mutated ubiquitin, our study revealed that only the overexpression of wild type and K6 ubiquitin (where all lysines except lysine 6 are mutated to arginine) resulted in reduced ubiquitination of CCT3^K21R^ compared to CCT3. This suggests that the ubiquitization at K21 of CCT3 is dominated by K6- linked ubiquitination (Fig 6C, S6C). Based on these experimental findings, we concluded that Sorafenib treatment leads to a decrease in K6-linked ubiquitination at K21 of CCT3. Furthermore, through an exogenous co-immunoprecipitation experiment in 293T cells, we observed that CCT3^K21R^ exhibited stronger binding to ACTN4 compared to CCT3 (Fig 6D, S6D). The results of molecular docking also showed a strong bond between CCT3^K21R^ and ACTN4 (Fig S6E). Importantly, the overexpression of both CCT3^K21R^ and CCT3 hindered the accumulation of ferrous ions in Sorafenib-treated 97L and LM3 cells (Fig 6E-F, S6F-G). Collectively, these results provide evidence that Sorafenib treatment leads to a reduction in K6- linked ubiquitination in K21 of CCT3, consequently facilitating its association with ACTN4 and its role in suppressing ferroptosis.

**Fig. 6.**
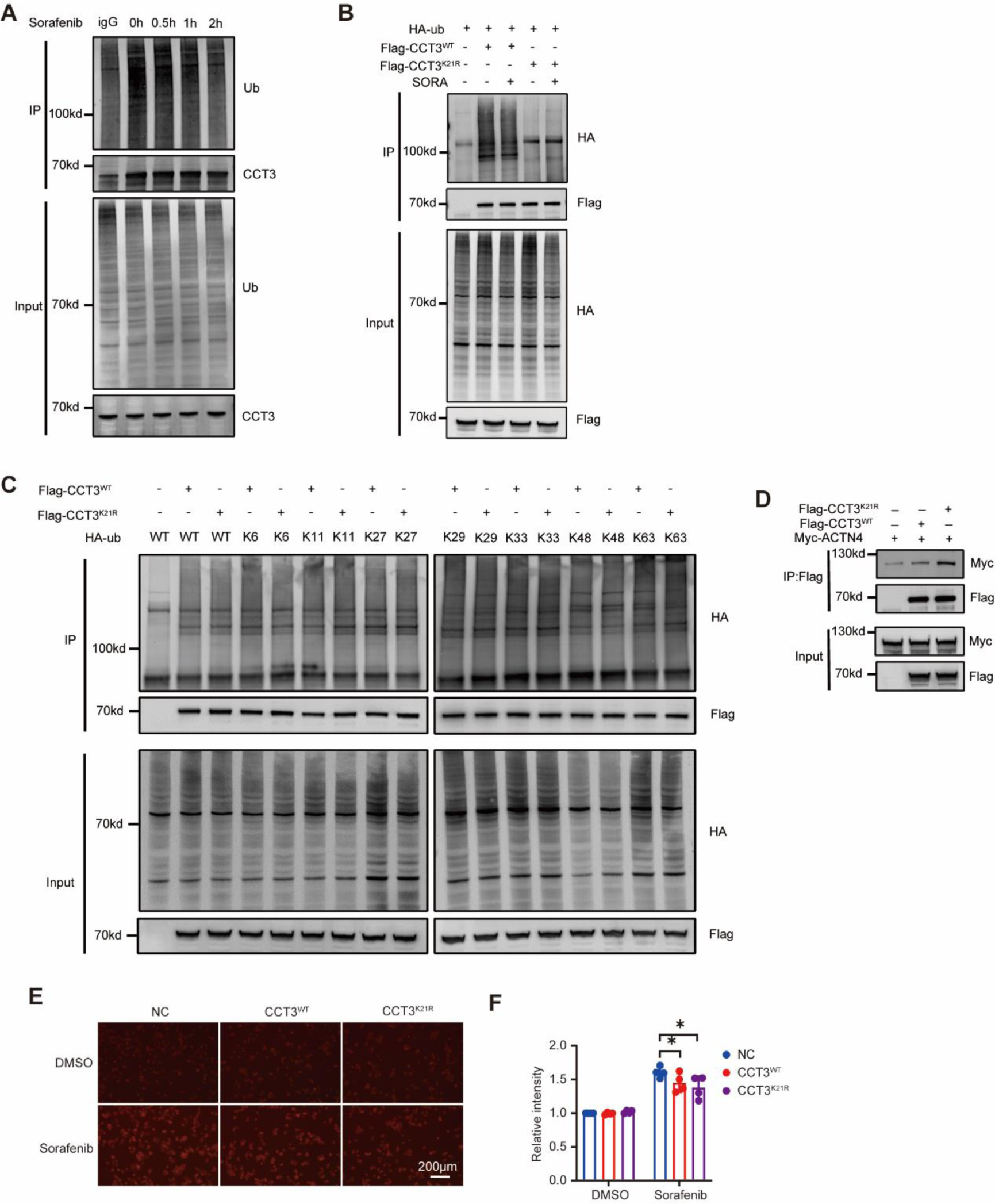
Ubiquitination at K21 was important for CCT3. A. The 97L cells were treated with Sorafenib (14 µM) for the indicated times and CCT3 ubiquitination was measured via immunoprecipitation-western blot assay. **B** After overexpression wild-type (CCT3^WT^) or K21R mutant CCT3 (CCT3^K21R^), its ubiquitination was measured in 293T cells treated with DMSO or Sorafenib (14µM) for 30 min. **C** 293T cells were transfected with plasmids expressing indicated wild-type or mutated proteins, CCT3 ubiquitination was measured via immunoprecipitation-western blot assay. K6, K11, K27, K29, K33, K48 and K63 indicates that all lysines except the indicated lysine have been mutated to arginine. **D** Detection of ACTN4 interactions with wild-type or K21R mutant CCT3 in 293T cells. **E-F** 97L cells overexpressed wild- type or K21R mutant CCT3 were treated with Sorafenib (14 µM) for 12 h, and then iron accumulation were measured. Histogram showing the statistical results from three biological replicates.

## Discussion

In 2013, Cong L et al. made the significant discovery that CRISPR arrays possess the capability to encode multiple guide sequences, thereby facilitating the simultaneous editing of multiple sites(Cong et al., 2013). As a result, the technique of CRISPR/Cas9 knockout library screening has undergone continuous enhancement and application in the identification of genes associated with the decline in cellular population fitness, encompassing diminished viability, heightened drug susceptibility, and decreased proliferation. Notably, the CRISPR/Cas9 knockout library screening method exhibits the capacity to manipulate nearly the entirety of the genome, encompassing non-coding elements. However, the limitations associated with off-target effects, as well as the presence of heterogeneous and heterozygote knockouts, undermine the dependability of its findings(Chan et al., 2022). PTM omics, on the other hand, presents a potent methodology for comprehensively characterizing PTMs at a system-wide level and has found extensive application in the identification of potential drug targets and the elucidation of drug mechanisms(Zhai et al., 2022). Nevertheless, it is important to note that while PTM omics can identify proteins implicated in drug action, it does not ascertain whether these proteins exert a positive or negative influence on the efficacy of the drug. Therefore, the integration of CRISPR/Cas9 knockout library screening and PTM omics serves to address the limitations of both techniques. Specifically, this study represents the first instance of combining CRISPR/Cas9 knockout library screening with PTM omics, leading to the identification of CCT3 as a factor that positively influences HCC resistance to Sorafenib. Furthermore, it was observed that the ubiquitination of CCT3 on K21 decreased following Sorafenib administration.

Our in vivo and in vitro experiments elucidate the underlying mechanism through which CCT3 contributes to Sorafenib resistance in HCC (Fig 7). Specifically, Sorafenib treatment diminishes the K6-linked ubiquitination of CCT3 on K21, consequently enhancing the interaction between CCT3 and ACTN4. This interaction, in turn, hampers the functionality of ACTN4 in facilitating the recycling of TFRC, thereby impeding the endocytotic TFRC from returning to the cell membrane. Consequently, this disruption leads to a reduction in the influx of iron ions into the cell. Consequently, it can be deduced that CCT3 exerts a negative regulatory influence on intracellular iron ion levels, ultimately inhibiting Sorafenib-induced ferroptosis.

**Fig. 7.**
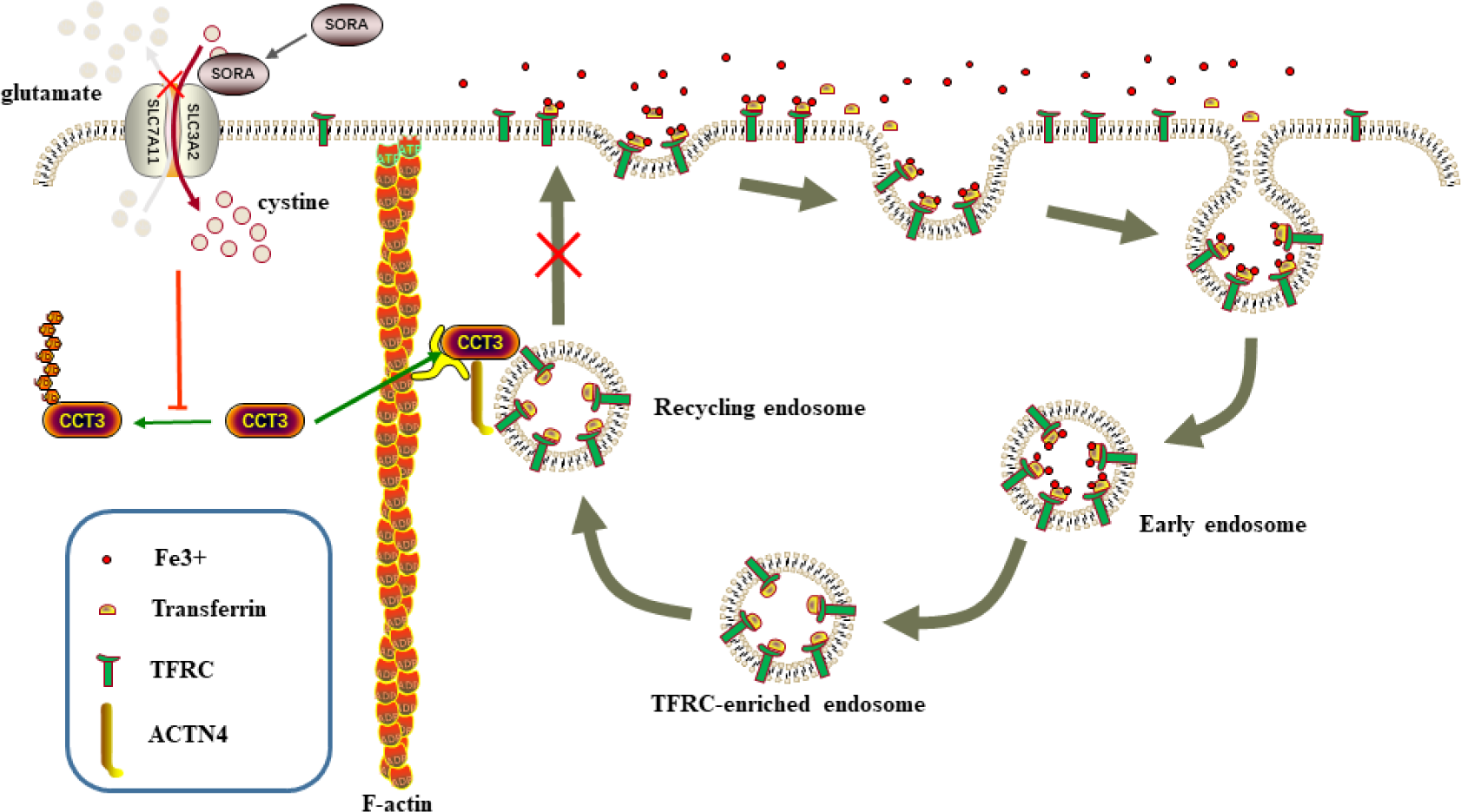
A model illustrating the function of CCT3 in inhibiting ferroptosis. CCT3 can inhibit the return of TFRC in TFRC-enriched endosome to the cell membrane by binding to ACTN4, which leads to a decrease in TFRC on the cell membrane and thus a decrease in iron ions transported into the cell, ultimately inhibiting ferroptosis. However, the K21 site of CCT3 is usually ubiquitinated so that it cannot perform its function of inhibiting ferroptosis, and its function is only maximized when sorafenib treatment results in the suppression of ubiquitination at this site.

CCT3 typically assumes a barrel-like conformation alongside the other seven members of the CCT family, wherein two rings of subunits enclose a central cavity(Vallin and Grantham, 2019). This multi-subunit oligomer plays a crucial role in the folding of cytoskeletal proteins and the maintenance of cellular proteostasis(Gestaut et al., 2019). Additionally, CCT3 can also function independently by interacting with YAP and TFCP2 in liver cancer, thereby preventing the PCBP2-induced ubiquitination of these proteins(Liu et al., 2019). In the present study, we observed that CCT3 has the ability to bind to ACTN4 and impede its functionality. However, due to the consistent expression of subunits within the CCT complex, altering the expression of one subunit will inevitably impact the expression of other subunits. Consequently, it is challenging to ascertain definitively whether the observed function is solely attributed to CCT3 or the entire CCT complex. Thus, additional investigations are warranted to address this issue(Kim and Choi, 2019; Sergeeva et al., 2019). Presently, there is a dearth of literature pertaining to the PTM of CCT3; however, other subunits within the CCT complex, such as CCT2, have been characterized(Abe et al., 2009). Our study elucidated that the ubiquitination of CCT3 at K21 resulted in a modification of its function and provided the initial identification of the specific ubiquitin chain type at this site. Additionally, we observed that Sorafenib treatment attenuated this ubiquitination modification, although the precise underlying mechanism necessitates further investigation. Previous reports have highlighted the impact of CCT3 on the prognosis of HCC, primarily focusing on its role in promoting proliferation and inhibiting apoptosis. In contrast, our study innovatively demonstrated that CCT3 can confer resistance to Sorafenib, the frontline chemotherapeutic agent for HCC.

Ferroptosis encompasses multiple metabolic pathways, namely iron metabolism, GSH metabolism, and lipid metabolism. Iron metabolism, in particular, encompasses the processes of iron absorption, storage, and utilization. This aspect of iron metabolism is not only implicated in ferroptosis but also plays a crucial role in normal physiological functions such as heme synthesis and DNA replication(Zhang et al., 2022). Cellular iron absorption occurs through two mechanisms: mediated by TFRC, which transports iron bound to transferrin, or facilitated by solute carrier family 39 member 14 (SLC39A14), which transports iron that is not bound to transferrin(Fang et al., 2023). The observation of TFRC overexpression in certain tumor cells and the inhibitory effect of TFRC deletion on erastin-induced ferroptosis have been documented(Yang and Stockwell, 2008; Zhao et al., 2022). Previous studies have primarily focused on the regulation of ferroptosis by TFRC through its differential expression, yet the present study found no significant alteration in TFRC expression following Sorafenib treatment(Chen et al., 2020). Considering the involvement of TFRC-mediated iron uptake via endosomal pathways, it was hypothesized that Sorafenib and CCT3 might impact this process(Rochette et al., 2022). Moreover, previous research has indicated that ACTN4, the specific protein targeted by CCT3 as identified in this investigation, plays a role in the process of endocytosis(Araki et al., 2000; Burton et al., 2021; Hara et al., 2007). Consequently, we proceeded to investigate the impact of CCT3 on the endocytosis of TFRC by conducting a series of experiments involving mechanisms and representations. Our findings unequivocally demonstrate that CCT3 hinders the recycling of endosomes containing TFRC.

In conclusion, our study employed PTM omics and CRISPR/Cas9 knockout library screening to systematically identify CCT3 as a significant gene associated with Sorafenib resistance. Notably, our findings demonstrate that CCT3 hinders TFRC recycling through its interaction with ACTN4, and this regulatory mechanism is governed by K6-linked ubiquitination on the K21 sites. This study holds theoretical significance by uncovering novel functions of chaperones and a previously unknown negative feedback mechanism of ferroptosis. Furthermore, this study holds significant clinical implications in the realm of guiding chemotherapy treatments for HCC and addressing the issue of Sorafenib resistance. One potential avenue to overcome this resistance involves the development of drugs that can effectively inhibit the interaction between CCT3 and ACTN4. Previous investigations have demonstrated that ferroptosis can hinder tumor immunity, resulting in paradoxical effects(Kim et al., 2022). Consequently, it is imperative to conduct further research to investigate the impact of CCT3 on the tumor immune microenvironment.

## Methods

### Reagents

Anti-CCT3 antibody (proteintech, 10571-1-AP), anti-TFRC antibody (proteintech, 10084-2-AP), anti-ubiquitin antibody (Cell Signaling Technology, 3936), anti-GPX4 antibody (proteintech, 14432-1-AP), anti-NCOA4 antibody (abcam, ab86707), anti-Myc antibody (proteintech, 16286- 1-AP), anti-ACTN4 antibody (proteintech, 19096-1-AP), anti-ACSL4 antibody (Santa Cruz Biotechnology, sc-271800), anti-Ferritin antibody (Santa Cruz Biotechnology, sc-376594), anti- SLC7A11 antibody (Cell Signaling Technology, 12691S), anti-Flag antibody (Bioss, bsm- 33346M), anti-GAPDH antibody (Fdbio, FD0063), anti-DMT1 antibody (proteintech, 20507-1- AP), anti-HA antibody (Santa Cruz Biotechnology, sc-7392), protein A/G (Santa Cruz Biotechnology, sc-2003), FITC-transferrin (Rockland Immunochemicals, 009-0234), BODIPY 581/591 C11 (ThermoFisher, D3861), FerroOrange (Dojindo, F374), Sorafenib (CSNpharm, CSN10381), Z-VAD-FMK (Selleck, S7023), Ferrostatin-1 (Selleck, S7243), Necrostatin-1 (Selleck, S8037), PTMScan Phospho-Tyrosine Rabbit mAb (P-Tyr-1000) Kit (Cell Signaling Technology, 8803), PTMScan Ubiquitin Remnant Motif (K-ε-GG) Kit (Cell Signaling Technology, 5562), PTMScan Acetyl-Lysine Motif [Ac-K] Kit (Cell Signaling Technology, 13416), GSH/GSSG detection assay (Beyotime, S0053), lipofectamine (ThermoFisher, 56532), Neofect (NEOFECT, TF20121201).

### Cell culture

The 97L and LM3 cell lines were procured from the Meisen Chinese Tissue Culture Collections (MeisenCTCC). These cell lines were cultivated in Dulbecco’s modified Eagle’s medium (DMEM), supplemented with 10% fetal bovine serum (FBS) and 1% penicillin/streptomycin, at a temperature of 37°C. To ensure their authenticity, we conducted short tandem repeat profiling on all cell lines and also tested them for mycoplasma contamination.

### Western blot

The cells were lysed using RIPA lysis buffer supplemented with a protease inhibitor cocktail and PR-619 on ice for a duration of 20 minutes. After centrifugation at 15,000g for 15 minutes at 4°C, the resulting supernatants were collected and individually mixed with 5 × sodium dodecyl sulfate (SDS) loading buffer. Subsequently, a 4-10 µL sample of protein was loaded onto 4-12% gradient Bis-Tris gels and transferred onto polyvinylidene difluoride membranes for blotting. After being blocked with a 1 × blocking solution at room temperature for one hour, the membranes were subjected to overnight incubation at 4 °C with various primary antibodies (diluted at 1:500-1:1000). Subsequently, the membranes were washed three times with 1 × tris buffered saline tween (TBST) and then incubated with HRP-conjugated secondary antibodies for two hours at room temperature. Following another three washes with 1 × TBST, the protein lanes were visualized using enhanced chemiluminescence and analyzed utilizing the Chemiluminescent Imaging System.

### Cell viability assay

Cell viability was assessed by employing the Cell Counting Kit-8 (CCK8) in accordance with the guidelines provided by the manufacturer. The cells were seeded into 96-well plates at a density ranging from 5-10 × 10^3^ cells per well and subsequently subjected to Sorafenib treatment. Following a duration of 24-48 hours, the cell culture medium was substituted with fresh DMEM supplemented with 10% CCK8 solutions, and the plates were reintroduced into a 37°C incubator for a period of 2 hours. The absorbance at 450 nm was then quantified using a spectrophotometer.

### Cell apoptosis assay

Cell apoptosis was assessed utilizing the Annexin V/propidium iodide (PI) apoptosis detection kit in accordance with the manufacturer’s guidelines. Both adherent and nonadherent cells in the supernatant were harvested and subjected to two washes with phosphate-buffered saline (PBS). Subsequently, a 100 µL cell suspension, resuspended at a density of 1 × 10^6^ cells/ml, was treated with 5 µL of Annexin V and PI staining solution. The resulting FITC-conjugated Annexin V and DNA-bound PI were then analyzed using flow cytometry.

### RNA interference and plasmid transfection

The transfection of siRNA and plasmid was conducted using lipofectamine 3000 or Neofect in accordance with the manufacturer’s protocols. Cells were passaged at the appropriate density 12 hours prior to transfection, and transfection reagents, along with siRNA or plasmid, were separately mixed with opti-MEM and allowed to incubate for 5 minutes. The two mixtures were then combined and left to incubate for an additional 15 minutes before being added to the medium. The efficacy of knockdown or overexpression was assessed 48 hours post-transfection through Western blot or qPCR analysis. Supplementary Table 3 contains a compilation of siRNA sequences.

### Immunoprecipitation analysis

The 97L, LM3, and 293FT cell lines were harvested and lysed at 4°C using 0.5% NP40 for CO-immunoprecipitation or RIPA lysis buffer for immunoprecipitation. Following centrifugation at 15,000g for 20 minutes at 4°C, a supernatant was obtained and incubated overnight at 4°C with gentle shaking, using either a specific antibody or an isotype control IgG. After the addition of protein A/G agarose beads, incubation was continued for an additional 2 hours. The beads were then washed five times with PBS before being either boiled in SDS loading buffer or eluted using 0.15% trifluoroacetic acid (TFA).

### Immunofluorescence analysis

In this study, 97L and LM3 cells were seeded and treated in 24-well plates with cell climbing slices. The cells in the cell climbing slices were fixed using a 4% polyformaldehyde solution, and a membrane-breaking solution was employed to disrupt the cell membrane. Subsequently, the cells were blocked for 15 minutes using a blocking solution and then incubated overnight at 4° with primary antibodies. After washing the cells three times with a wash buffer, they were incubated with Alexa Fluor 555 and Alexa Fluor 488 conjugated secondary antibodies for 1 hour at room temperature. Finally, DAPI anti-quenching sealant was applied to the slides, and confocal laser microscope imaging was used to capture the images.

### Transferrin endocytosis assay

The cells in a 12-well plate were subjected to two washes with serum-free DMEM and subsequently incubated with the same medium at a temperature of 37°C for a duration of 1 hour. Following this, a serum-free DMEM medium containing 25 ug/ml of FITC-transferrin was introduced, and the cells were further incubated at 37°C for a period of 15 minutes. The adherent cells were dissociated using 0.25% trypsin-EDTA, fixed with 4% paraformaldehyde at room temperature for 20 minutes, and subjected to two washes with PBS. The intracellular FITC-transferrin signals were then quantified using a flow cytometer equipped with FITC detection settings.

### GSH/GSSG detection assay

The intracellular levels of glutathione (GSH) and oxidized glutathione (GSSG) were assessed using GSH/GSSG assay kits following the manufacturer’s instructions. In summary, cells in a 24-well plate were harvested and homogenized with protein removal agent M solution. Subsequently, the samples underwent two cycles of freezing and thawing using liquid nitrogen and 37°C water baths. After a 5-minute incubation on ice, the supernatant was obtained by centrifugation at 10,000g for 10 minutes for the quantification of glutathione and oxidized glutathione.

### Transmission electron microscopy

The cells were subjected to fixation for a duration of 24 hours using a 2.5% glutaraldehyde solution. Following three washes with PBS, the cells were exposed to a 1% osmic acid solution for 1 hour, followed by treatment with a 2% uranium acetate solution for 30 minutes. Subsequently, the sample underwent dehydration and embedding processes, enabling the creation of ultrathin sections. Digital images were obtained using transmission electron microscopy at magnifications of 2000× and 8500×.

### Colony formation assay

The LM3 and 97L cells were suspended in a solution of 10% FBS and DMEM. A seeding density of 1000 cells per well was utilized in six-well plates, followed by treatment with Sorafenib at the specified concentrations. The cells were maintained in a humidified incubator at a temperature of 37°, and the medium was replaced every 2 to 3 days. After a duration of two weeks, the cells were fixed using a 4% solution of paraformaldehyde for a period of 20 minutes. Subsequently, the colonies were stained with a 0.1% solution of crystal violet for 20 minutes and washed with PBS three times. The colonies were then photographed and quantified using ImageJ software.

### Iron staining

Intracellular labile iron was assessed utilizing the metal sensor FerroOrange. In brief, a total of approximately 5 x 10^4^ cells were seeded per well in a 24-well plate, and the designated treatments were administered to the cells on the subsequent day. Following a 12-hour incubation period, FerroOrange was introduced at a dilution of 2500 times and incubated at a temperature of 37°C for a duration of 10 minutes, after which it was captured via fluorescence microscopy. The estimation of intracellular labile iron was determined based on the intensity of yellow fluorescence and quantified using imageJ software.

### Lipid ROS assay

Intracellular non-peroxidized lipids were assessed by employing the BODIPY 581/591 C11 dye. To initiate the experiment, approximately 1 x 10^5^ cells were seeded per well in a 12-well plate, and the designated treatments were administered to the cells on the subsequent day. Following a 24-hour incubation period, BODIPY 581/591 C11 was introduced and allowed to incubate at a temperature of 37°C for a duration of 15 minutes. Subsequently, the cells were fixed with a 4% paraformaldehyde solution, and the non-peroxidized lipids were evaluated using flow cytometry.

### Molecular docking

The crystal structures of TCPG (7LUM) and ACTN4 (6OA6) were obtained from the Protein Data Bank(Feng et al., 2020; Knowlton et al., 2021). The protein was prepared using AutoDockTools-1.5.7, with manual removal of water molecules and addition of polar hydrogen(Morris et al., 2008). Protein-protein docking was conducted using the Docking Web Server (GRAMM)(Katchalski-Katzir et al., 1992; Vakser, 1996). The resulting protein-protein complex underwent manual optimization by AutoDockTools-1.5.7 to eliminate water and add polar hydrogen. Finally, PyMOL was utilized to predict the protein-protein interactions and generate the corresponding figure.

### Protein extraction and digestion

The 97L and LM3 cells were harvested and lysed in freshly prepared lysis buffers containing 8 M urea, 150 mM NaCl, 50 mM Tris-HCl pH 8.0, 1 × phosphorylase inhibitor, 1 × deacetylase inhibitor, 50 mM (2,6-diamino-5-thiocyanatopyridin-3-yl) thiocyanate (PR- 619), and 1× protease inhibitor cocktail at a temperature of 4°C for a duration of 20 minutes. The protein concentration of the supernatant was estimated using the bicinchoninic acid protein assay (BCA). Protein reduction was achieved by treating with dithiothreitol for 45 minutes at a temperature of 30°C, followed by carbamidomethylation using Iodoacetamide for 30 minutes at room temperature. Subsequently, the solvent was substituted with 50 mmol/L ammonium bicarbonate using a Zeba Spin Desalting Column or Ultrafiltration centrifugal tube. The samples were subjected to overnight digestion at 37°C, employing a trypsin-to-substrate ratio of 1:40 (wt/wt) and agitation at 150 rpm. The peptides were acidified with trifluoroacetic acid (TFA), desalted through C18 Sep-Pak SPE cartridges, and subsequently subjected to vacuum drying.

### Enrichment of PTM peptides

Proteomics analyses were conducted to investigate acetylation, phosphorylation, and ubiquitination. Digestive peptides were resuspended in ice-cold 1 × IAP buffer and incubated with cross-linked antibody beads for a duration of two hours at 4°C. The elution of peptides was achieved using 0.15% TFA after three rounds of soft-washing with ice-cold 1 × IAP buffer and two rounds with ice-cold ultrapure water. The eluate was completely dried through vacuum centrifugation and subsequently utilized for subsequent analysis.

### Tandem mass tag (TMT) isobaric labeling

The TMT Label Reagents were equilibrated at room temperature prior to use and subsequently dissolved in anhydrous acetonitrile for a duration of 5 minutes, with intermittent vortexing. A peptide sample was dissolved in a solution containing 100 mmol/L of tetraethylammonium bromide at a pH of 8.0. Following this, the TMT Label Reagent was added to the peptide sample and incubated at room temperature for a period of one hour. To halt the reaction, all samples were treated with 5% hydroxylamine for 15 minutes and combined into a single tube. The TMT-labeled peptides were then dried using vacuum centrifugation after desalting with a C18 stage tip that was prepared in-house.

### Strong anion exchange (SAX) fractionation

Empore Anion-SR Extraction Disks were utilized for the purpose of performing SAX chromatography using tip columns. The SAX tip, which was prepared in-house, underwent conditioning and equilibration with 100 μL of acetonitrile (ACN), followed by 100 μL of SAX B, 200 μL of SAX C, and finally 100 μL of SAX A. Mixtures containing PTM peptides labeled with TMT were reconstituted in 100 μL of SAX A and subsequently loaded onto the previously treated SAX tips. Upon loading the SAX tips with 100 μL of SAX buffers 1, 2, 3, 4, and 5, the resulting eluates were designated as SAX fractions 1, 2, 3, 4, and 5, respectively. Supplementary Table 4 provides a comprehensive list of the formulations for the SAX reagents.

### Liquid chromatography–tandem mass spectrometry (LC-MS/MS) analysis

The peptide samples obtained after SAX fractionation underwent desalting and vacuum evaporation, followed by resuspension in a solution containing 2% ACN and 0.1% formic acid (FA). LC-MS/MS analysis was conducted using a Thermo Fisher Scientific Orbitrap Exploris 480 mass spectrometer coupled to an UltiMate NCS-3500RSC system. Gradient elution was performed at a flow rate of 450 nL/min, with linear gradients lasting for a total of 120 minutes. Specifically, the solvent composition was as follows: 3%-10% solvent B from 1 to 5 minutes, 10%-26% solvent B from 6 to 90 minutes, 26%-38% solvent B from 91 to 110 minutes, 38%-80% solvent B from 111 to 115 minutes, and 80% solvent B from 116 to 120 minutes. The MS spectra were obtained with a resolution of 60,000, covering a mass range of 400–1200 m/z, and utilizing a normalized automatic gain control (AGC) target of 300%. For the acquisition of MS2 spectra, a resolution of 15,000 was employed, along with a higher energy collision induced dissociation (HCD) collision energy set at 35%. The isolation window was set at 1.6 m/z, while the dynamic exclusion window lasted for a duration of 30 seconds.

### Protein identification and quantification

The identification and quantification of proteins were conducted using MaxQuant (version 1.6.2.10). The MS2-based TMT18plex quantification was employed for the whole cell proteome, while the MS2-based TMT10plex quantification was utilized for the acetylome, phosphoproteome, and ubiquitinome. The human UniProtKB served as the search database, with the automatic reverse database and known contaminants employed as decoys. The carbamidomethyl-cysteine modification was designated as a fixed modification, while the N-terminal acetylation and methionine oxidation modifications were considered as variable modifications. In addition, the variable modifications encompassing the acetylome, phosphoproteome, and ubiquitinome were incorporated. The upper limit for modifications per peptide was established at five. The specific enzyme employed was trypsin, with a tolerance of two missed cleavages per peptide. Any parameters not explicitly mentioned were assigned the default settings of Maxquant.

### Bioinformatics analysis

The functional annotation enrichment analysis for proteins with altered PTM or expression was conducted using the clusterProfiler package in R(Yu et al., 2012). The FerrDb database was utilized to obtain the Ferroptosis related pathway in the GSEA analysis(Zhou et al., 2023). Additionally, the consensus motifs of the PTM were examined by analyzing sequences within - 15 and +15 amino acids of the PTM sites, and the resulting amino acid sequence diagram was generated using the ggseqlogo package in R.

### Statistical analysis

The statistical and bioinformatics data were predominantly analyzed using the R framework (version 4.3.0) and GraphPad Prism 9. Unless specified otherwise, the experiments were conducted independently in triplicates, and the data is presented as mean ± SD. All statistical significance was determined using a two-tailed, Student’s t-test, unless otherwise stated. A *P*- value of less than 0.05 was considered statistically significant (^ns^*P* ≥ 0.05; ^✱^*P* < 0.05, ^✱✱^*P* < 0.01).

### Data availability

The mass spectrometry proteomics data deposited to the ProteomeXchange Consortium via the iProX partner repository has been assigned the dataset identifier PXD045576(Chen et al., 2022b; Ma et al., 2019). Supplementary information containing additional data supporting our findings is also provided.

## Acknowledgements

This study has received support from the National Key Research and Development Program of China (grant number 2021YFC2301805), the National Natural Science Foundation of China (grant numbers 82203610, 81972597), and the opening foundation of the State Key Laboratory for Diagnosis and Treatment of Infectious Diseases (grant number SKLID2023KF06).

## Author contributions

H.Z., Q.L., Q.M., J.L., N. A., L.J., X.O., and X.L. performed the experiments. D.Z., Y.C., and S.Z. performed the majority of data and statistical analysis. F.J. and L.Z. conceived and designed experiments. F.J. and Q.M. wrote and edited the manuscript. F.J., L.W., and L.L. directed the study.

## Competing interests

The authors declare no competing interests.

---

**Fig. S1.**
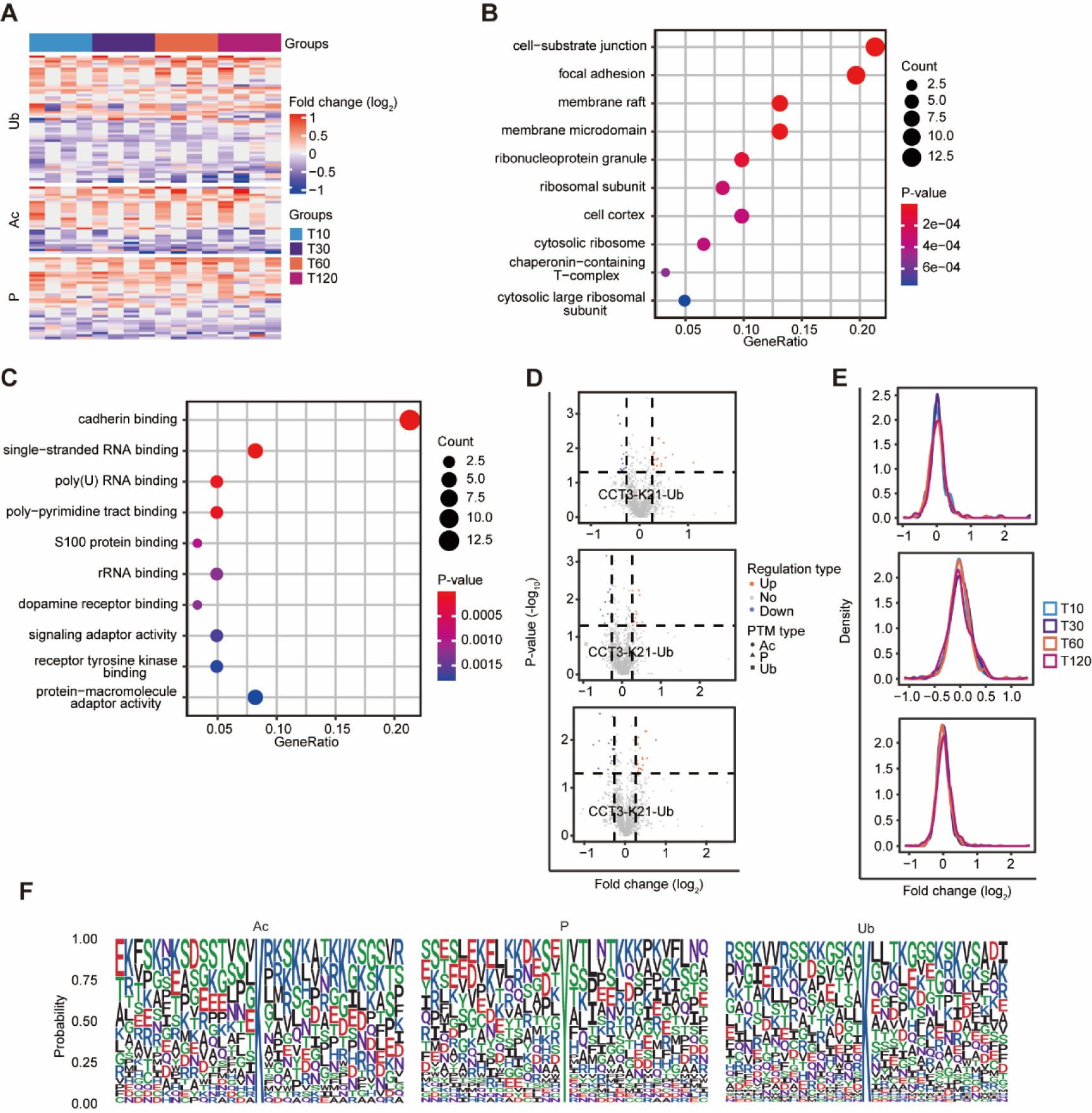
PTM omics for 97L cells under Sorafenib treatment. A. Heat maps showed PTM sites with significant regulated in ubiquitination, acetylation and phosphorylation in 97L cells after Sorafenib treatment. **B-C** GO enrichment analysis (cell component and molecular function) of proteins containing regulated PTM site after Sorafenib treatment. **D** Volcano plot showing regulated PTM sites in 97L cells treated with Sorafenib for 10 min, 30 min, and 2 h. **E** Density distribution map of acetylation, phosphorylation and ubiquitination sites after Sorafenib treatment. “T10” means “10 min vs 0 min”. **F** iceLogo plots showing the difference of amino acid frequency at positions flanking the Sorafenib-regulated acetylation, phosphorylation and ubiquitination sites.

**Fig. S2.**
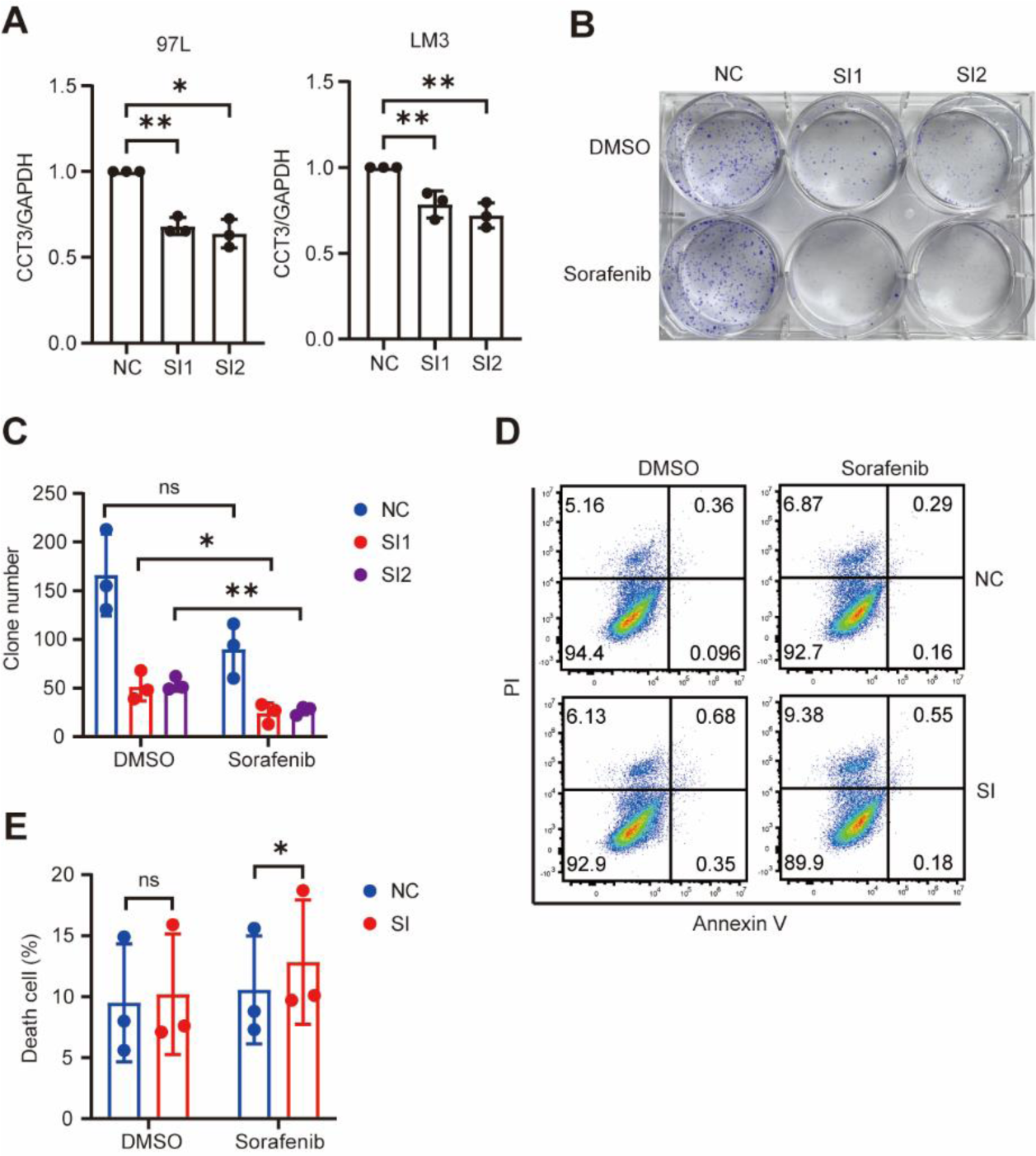
The inhibition of CCT3 sensitized HCC cells to Sorafenib. A. The expression of CCT3 in indicted groups detected by Western blot were quantified. Statistical analysis was performed from three biological replicates. **B-C** Clone formation capacity of control and CCT3-knockdown LM3 cells following treatment with DMSO or 2 µM Sorafenib. Statistical analysis was performed from three biological replicates. **D** Apoptosis of indicted LM3 cells treated with DMSO or 7 µM Sorafenib for 24 h were performed using Annexin V/PI staining. **E** Histogram showing the statistical results of apoptosis from three biological replicates.

**Fig. S3.**
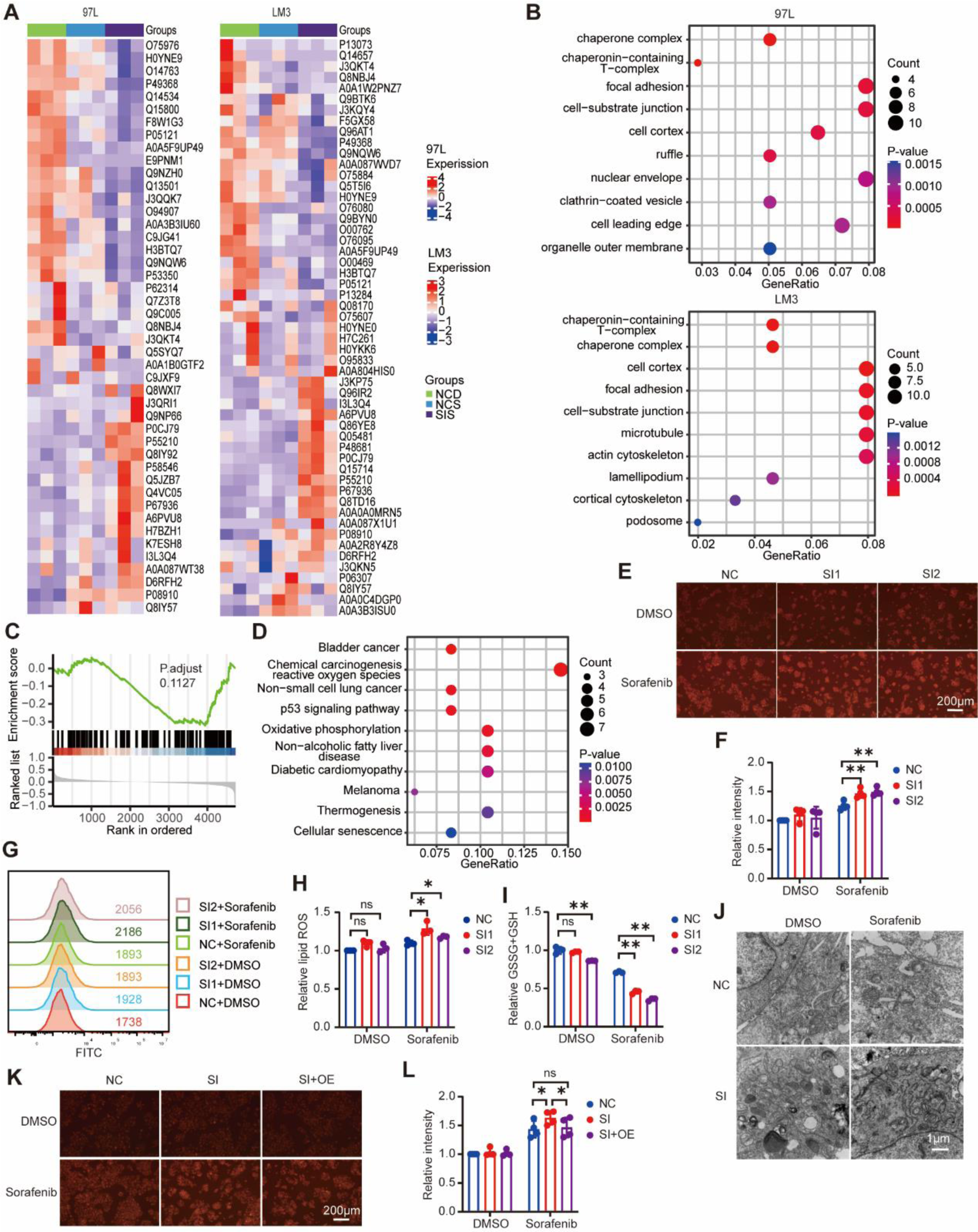
The inhibition of CCT3 promotes ferroptosis in HCC cells treated with Sorafenib. A. Heat map showing the differential gene expression between different groups (NCD: control + DMSO, NCS: control + Sorafenib, SIS: CCT3 knockdown + Sorafenib) in indicated cells. **B** GO enrichment analysis (cell component) of differentially expressed protein between control and CCT3-knockdown cells. **C** GSEA ferroptosis pathway enrichment analysis of differentially expressed protein between DMSO and Sorafenib treatment 97L cells. **D** KEGG pathway enrichment analysis of differentially expressed protein between DMSO and Sorafenib treatment LM3 cells. **E-F** The accumulation of iron in control and CCT3-knockdown LM3 cells were detected using FerroOrange after treated with DMSO or Sorafenib (14 µM) for 12 h. Statistical results of iron accumulation were from four biological replicates. **G-H** LM3 cells were treated with DMSO or Sorafenib (14 µM) for 24 h, and then lipid hydroperoxides were measured. Statistics for the median Fluorescence intensity of oxidized BODIPY dyes were carried out on three biological replicates. **I** LM3 cells were treated with Sorafenib at 14 µM for 12 h, and the intracellular glutathione (GSH) level were assayed. **J** Transmission electron microscopy was used to examine the morphology of indicated LM3 cells after 24 hours of treatment with Sorafenib (14 µM). **K-L** Indicated LM3 cells were treated with Sorafenib (14 µM) for 12 h, and then iron accumulation were measured and statistical analysis.

**Fig. S4.**
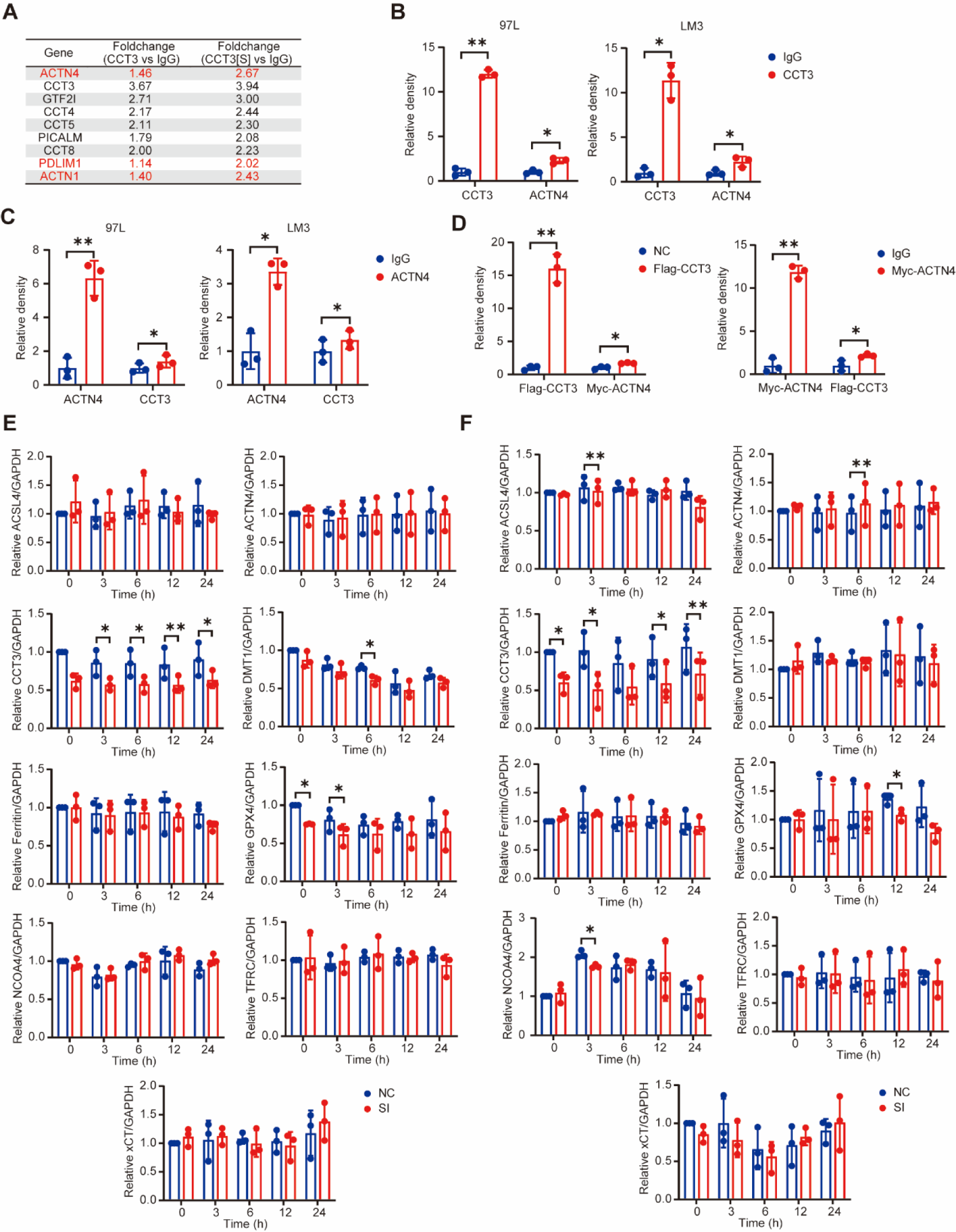
An interacting protein of CCT3 is ACTN4. A. List of proteins with significant changes between indicated groups. **B** Quantitative and statistical analysis of western blot result of CCT3-binding protein in 97L and LM3 cells. **C** Quantitative and statistical analysis of Western blot result of ACTN4-binding protein in 97L and LM3 cells. **D** Quantitative and statistical analysis of western blot result of CCT3-binding and ACTN4-binding protein in 293T cells. **E-F** The expression of CCT3, ACTN4 and protein associated with ferroptosis in indicted groups detected by Western blot were quantified and analyzed. ALL statistical analysis was performed from three biological replicates.

**Fig. S5.**
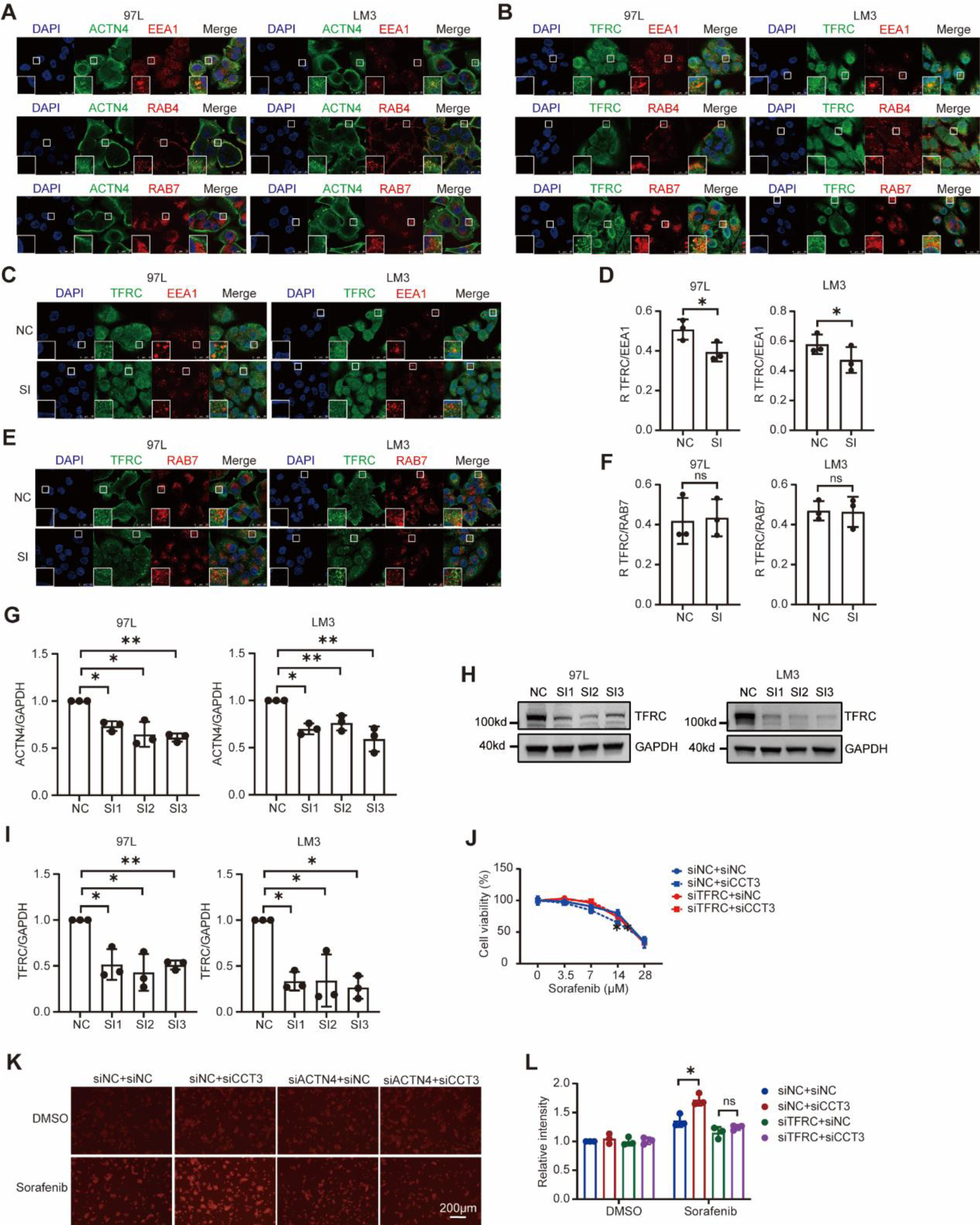
Iron endocytosis is inhibited by CCT3 by impairing TFRC recycling through ACTN4. A. Immunofluorescence staining showing the distribution of ACTN4 (green), EEA1 (red), RAB4 (red), RAB7 (red) and DAPI (blue) in indicated cells. **B** Immunofluorescence staining showing the distribution of TFRC (green), EEA1 (red), RAB4 (red), RAB7 (red) and DAPI (blue) in indicated cells. **C-F** The distribution of EEA1 (red), RAB7 (red), TFRC (green) and DAPI (blue) in control and CCT3-knockdown cells. Histogram showing the statistical results for Pearson’s R correlation value from three biological replicates. **G** The expression of ACTN4 in indicted cells detected by Western blot were quantified and analyzed. **H-I** Western blot analysis of the knockdown efficiency of TFRC in 97L and LM3 cells. Histogram showing the statistical results from three biological replicates. **J** Cell viability analysis of control, CCT3- knockdown, and ACTN4-knockdown LM3 cells following treatment with Sorafenib for 24 h. **K-L** Indicated LM3 cells were treated with Sorafenib (14 µM) for 12 h, and then iron accumulation were measured. Histogram showing the statistical results from three biological replicates.

**Fig. S6.**
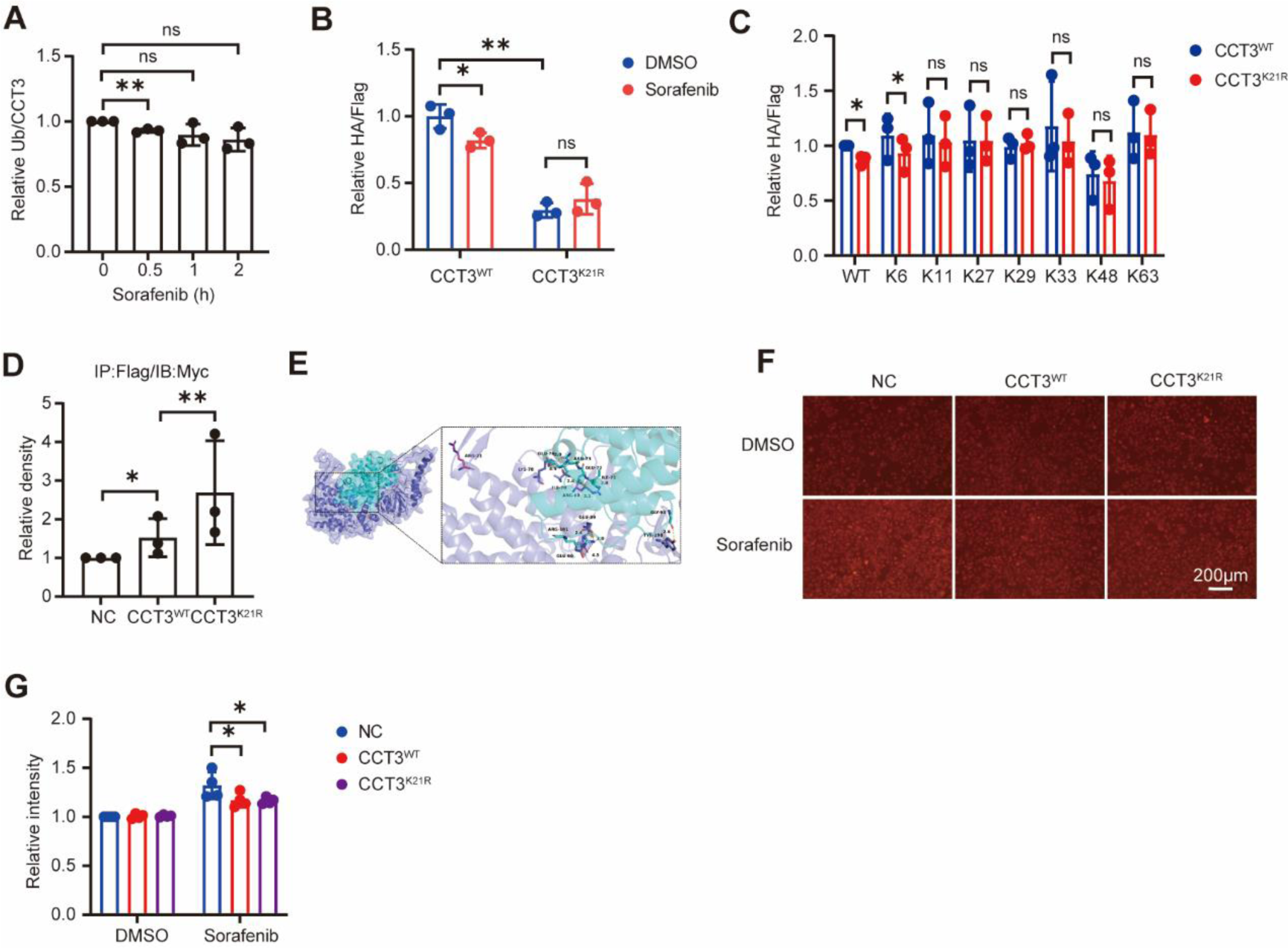
K21 ubiquitination was essential for CCT3. A. The ubiquitinated CCT3 after Sorafenib treatment was quantified and analyzed from three biological replicates in 97L cells. **B** The ubiquitination of wild-type or K21R mutant CCT3 after Sorafenib treatment was quantified and analyzed from three biological replicates in 293T cells. **C** Ubiquitination levels of exogenous CCT3 measured by immunoprecipitation-western blot assay was quantified and analyzed from three biological replicates in 293T cells. **D** Quantification and statistics of ACTN4 interactions with wild-type or K21R mutant CCT3 in 293T cells. **E** Molecular docking of the CCT3^K21R^ (slate cartoon) and ACTN4(cyan cartoon) interaction complex, with corresponding-colored stick structures representing the binding sites. **F-G** LM3 cells overexpressed wild-type or K21R mutant CCT3 were treated with Sorafenib (14 µM) for 12 h, and then iron accumulation were measured. Histogram showing the statistical results from three biological replicates.

## Notes

### Competing Interest Statement

The authors have declared no competing interest.

